# Optimizer’s dilemma: optimization strongly influences model selection in transcriptomic prediction

**DOI:** 10.1101/2023.06.26.546586

**Authors:** Jake Crawford, Maria Chikina, Casey S. Greene

## Abstract

**Motivation:** Most models can be fit to data using various optimization approaches. While model choice is frequently reported in machine-learning-based research, optimizers are not often noted. We applied two different implementations of LASSO logistic regression implemented in Python’s scikit-learn package, using two different optimization approaches (coordinate descent and stochastic gradient descent), to predict driver mutation presence or absence from gene expression across 84 pan-cancer driver genes. Across varying levels of regularization, we compared performance and model sparsity between optimizers.

**Results:** After model selection and tuning, we found that coordinate descent (implemented in the `liblinear` library) and SGD tended to perform comparably. `liblinear` models required more extensive tuning of regularization strength, performing best for high model sparsities (more nonzero coefficients), but did not require selection of a learning rate parameter. SGD models required tuning of the learning rate to perform well, but generally performed more robustly across different model sparsities as regularization strength decreased. Given these tradeoffs, we believe that the choice of optimizers should be clearly reported as a part of the model selection and validation process, to allow readers and reviewers to better understand the context in which results have been generated.

**Availability and implementation:** The code used to carry out the analyses in this study is available at https://github.com/greenelab/pancancer-evaluation/tree/master/01_stratified_classification. Performance/regularization strength curves for all genes in the Vogelstein et al. 2013 dataset are available at https://doi.org/10.6084/m9.figshare.22728644.

## Introduction

Gene expression profiles are widely used to classify samples or patients into relevant groups or categories, both preclinically [1,2] and clinically [3,4]. To extract informative gene features and to perform classification, a diverse array of algorithms exist, and different algorithms perform well across varying datasets and tasks [1]. Even within a given model class, multiple optimization methods can often be applied to find well-performing model parameters or to optimize a model’s loss function. One commonly used example is logistic regression. The widely used scikit-learn Python package for machine learning [5] provides two modules for fitting logistic regression classifiers: `LogisticRegression`, which uses the `liblinear` coordinate descent method [6] to find parameters that optimize the logistic loss function, and `SGDClassifier`, which uses stochastic gradient descent [7] to optimize the same loss function.

Using scikit-learn, we compared the `liblinear` (coordinate descent) and SGD optimization techniques for prediction of driver mutation status in tumor samples, across a wide variety of genes implicated in cancer initiation and development [8]. We applied LASSO (L1-regularized) logistic regression, and tuned the strength of the regularization to compare model selection between optimizers. We found that across a variety of models (i.e. varying regularization strengths), the training dynamics of the optimizers were considerably different: models fit using `liblinear` tended to perform best at fairly high regularization strengths (100-1000 nonzero features in the model) and overfit easily with low regularization strengths. On the other hand, after tuning the learning rate, models fit using SGD tended to perform well across both higher and lower regularization strengths, and overfitting was less common.

Our results caution against viewing optimizer choice as a “black box” component of machine learning modeling. The observation that LASSO logistic regression models fit using SGD tended to perform well for low levels of regularization, across diverse driver genes, runs counter to conventional wisdom in machine learning for high-dimensional data which generally states that explicit regularization and/or feature selection is necessary. Comparing optimizers or model implementations directly is rare in applications of machine learning for genomics, and our work shows that this choice can affect generalization and interpretation properties of the model significantly. Based on our results, we recommend considering the appropriate optimization approach carefully based on the goals of each individual analysis.

## Methods

### Data download and preprocessing

To generate binary mutated/non-mutated gene labels for our machine learning model, we used mutation calls for TCGA Pan-Cancer Atlas samples from MC3 [9] and copy number threshold calls from GISTIC2.0 [10]. MC3 mutation calls were downloaded from the Genomic Data Commons (GDC) of the National Cancer Institute, at https://gdc.cancer.gov/about-data/publications/pancanatlas.

Thresholded copy number calls are from an older version of the GDC data and are available here: https://figshare.com/articles/dataset/TCGA_PanCanAtlas_Copy_Number_Data/6144122. We removed hypermutated samples, defined as two or more standard deviations above the mean non-silent somatic mutation count, from our dataset to reduce the number of false positives (i.e., non-driver mutations). Any sample with either a non-silent somatic variant or a copy number variation (copy number gain in the target gene for oncogenes and copy number loss in the target gene for tumor suppressor genes) was included in the positive set; all remaining samples were considered negative for mutation in the target gene.

RNA sequencing data for TCGA was downloaded from GDC at the same link provided above for the Pan-Cancer Atlas. We discarded non-protein-coding genes and genes that failed to map, and removed tumors that were measured from multiple sites. After filtering to remove hypermutated samples and taking the intersection of samples with both mutation and gene expression data, 9074 total TCGA samples remained.

### Cancer gene set construction

In order to study mutation status classification for a diverse set of cancer driver genes, we started with the set of 125 frequently altered genes from Vogelstein et al. [8] (all genes from Table S2A). For each target gene, in order to ensure that the training dataset was reasonably balanced (i.e., that there would be enough mutated samples to train an effective classifier), we included only cancer types with at least 15 mutated samples and at least 5% mutated samples, which we refer to here as “valid” cancer types. In some cases, this resulted in genes with no valid cancer types, which we dropped from the analysis. Out of the 125 genes originally listed in the Vogelstein et al. cancer gene set, we retained 84 target genes after filtering for valid cancer types.

### Classifier setup and optimizer comparison details

We trained logistic regression classifiers to predict whether or not a given sample had a mutational event in a given target gene using gene expression features as explanatory variables. Based on our previous work, gene expression is generally effective for this problem across many target genes, although other -omics types can be equally effective in many cases [11]. Our model was trained on gene expression data (X) to predict mutation presence or absence (y) in a target gene. To control for varying mutation burden per sample and to adjust for potential cancer type-specific expression patterns, we included one-hot encoded cancer type and log10(sample mutation count) in the model as covariates. Since gene expression datasets tend to have many dimensions and comparatively few samples, we used a LASSO penalty to perform feature selection [12]. LASSO logistic regression has the advantage of generating sparse models (some or most coefficients are 0), as well as having a single tunable hyperparameter which can be easily interpreted as an indicator of regularization strength, or model complexity.

To compare model selection across optimizers, we first split the “valid” cancer types into train (75%) and test (25%) sets. We then split the training data into “subtrain” (66% of the training set) data to train the model on, and “holdout” (33% of the training set) data to perform model selection, i.e. to use to select the best-performing regularization parameter, and the best-performing learning rate for SGD in the cases where multiple learning rates were considered. In each case, these splits were stratified by cancer type, i.e. each split had as close as possible to equal proportions of each cancer type included in the dataset for the given driver gene.

### LASSO parameter range selection and comparison between optimizers

The scikit-learn implementations of coordinate descent (in `liblinear` / `LogisticRegression`) and stochastic gradient descent (in `SGDClassifier`) use slightly different parameterizations of the LASSO regularization strength parameter. `liblinear` ‘s logistic regression solver optimizes the following loss function:

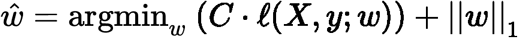

where 𝓁(*X, y*; *w*) denotes the negative log-likelihood of the observed data (*X, y*) given a particular choice of feature weights *w*. `SGDClassifier` optimizes the following loss function:

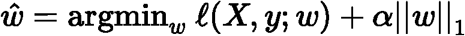

which is equivalent with the exception of the LASSO parameter which is formulated slightly differently, as 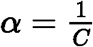. The result of this slight difference in parameterization is that `liblinear` *C* values vary inversely with regularization strength (higher values = less regularization, or greater model complexity) and `SGDClassifier` *α* values vary directly with regularization strength (lower values = less regularization, or greater model complexity).

For the `liblinear` optimizer, we trained models using *C* values evenly spaced on a logarithmic scale between (10^−3^, 10^7^); i.e. the output of `numpy.logspace(−3, 7, 21)`. For the SGD optimizer, we trained models using the inverse range of *α* values between (10^−7^, 10^3^), or `numpy.logspace(−7, 3, 21)`. These hyperparameter ranges were intended to give evenly distributed coverage across genes that included “underfit” models (predicting only the mean or using very few features, poor performance on all datasets), “overfit” models (performing perfectly on training data but comparatively poorly on cross-validation and test data), and a wide variety of models in between that typically included the best fits to the cross-validation and test data.

For ease of visual comparison in our figures, we plot the SGD *α* parameter directly, and the `liblinear` *C* parameter inversely (i.e. 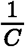). This orients the x-axes of the relevant plots in the same direction: lower values represent lower regularization strength or higher model complexity, and higher values represent higher regularization strength or lower model complexity, for both optimizers.

### SGD learning rate selection

scikit-learn’s `SGDClassifier` provides four built-in approaches to learning rate scheduling: `constant` (a single, constant learning rate), `optimal` (a learning rate with an initial value selected using a heuristic based on the regularization parameter and the data loss, that decreases across epochs), `invscaling` (a learning rate that decreases exponentially by epoch), and `adaptive` (a learning rate that starts at a constant value, which is divided by 5 each time the training loss fails to decrease for 5 straight epochs). The `optimal` learning rate schedule is used by default.

When we compared these four approaches, we used a constant learning rate of 0.0005, and an initial learning rate of 0.1 for the `adaptive` and `invscaling` schedules. We also tested a fifth approach that we called “ `constant_search` “, in which we tested a range of constant learning rates in a grid search on a validation dataset, then evaluated the model on the test data using the best-performing constant learning rate by validation AUPR. For the grid search, we used the following range of constant learning rates: {0.000005, 0.00001, 0.00005, 0.0001, 0.0005, 0.001, 0.01}. Unless otherwise specified, results for SGD in the main paper figures used the `constant_search` approach, which performed the best in our comparison between schedulers.

## Results

### liblinear and SGD LASSO models perform comparably, but liblinear is sensitive to regularization strength

For each of the 125 driver genes from the Vogelstein et al. 2013 paper, we trained models to predict mutation status (presence or absence) from RNA-seq data, derived from the TCGA Pan-Cancer Atlas. For each optimizer, we trained LASSO logistic regression models across a variety of regularization parameters (see Methods for parameter range details), achieving a variety of different levels of model sparsity (Supplementary Figure S1). We repeated model fitting/evaluation across 4 cross-validation splits x 2 replicates (random seeds) for a total of 8 different models per parameter. Cross-validation splits were stratified by cancer type.

Previous work has shown that pan-cancer classifiers of Ras mutation status are accurate and biologically informative [13]. We first evaluated models for KRAS mutation prediction. As model complexity increases (more nonzero coefficients) for the `liblinear` optimizer, we observed that performance increases then decreases, corresponding to overfitting for high model complexities/numbers of nonzero coefficients (Figure 1A). On the other hand, for the SGD optimizer, we observed consistent performance as model complexity increases, with models having no nonzero coefficients performing comparably to the best (Figure 1B). In this case, top performance for SGD (a regularization parameter of 10^−1^) is slightly better than top performance for `liblinear` (a regularization parameter of 1 / 3.16 × 10^2^): we observed a mean test AUPR of 0.722 for SGD vs. mean AUPR of 0.692 for `liblinear`.

**Figure 1:**
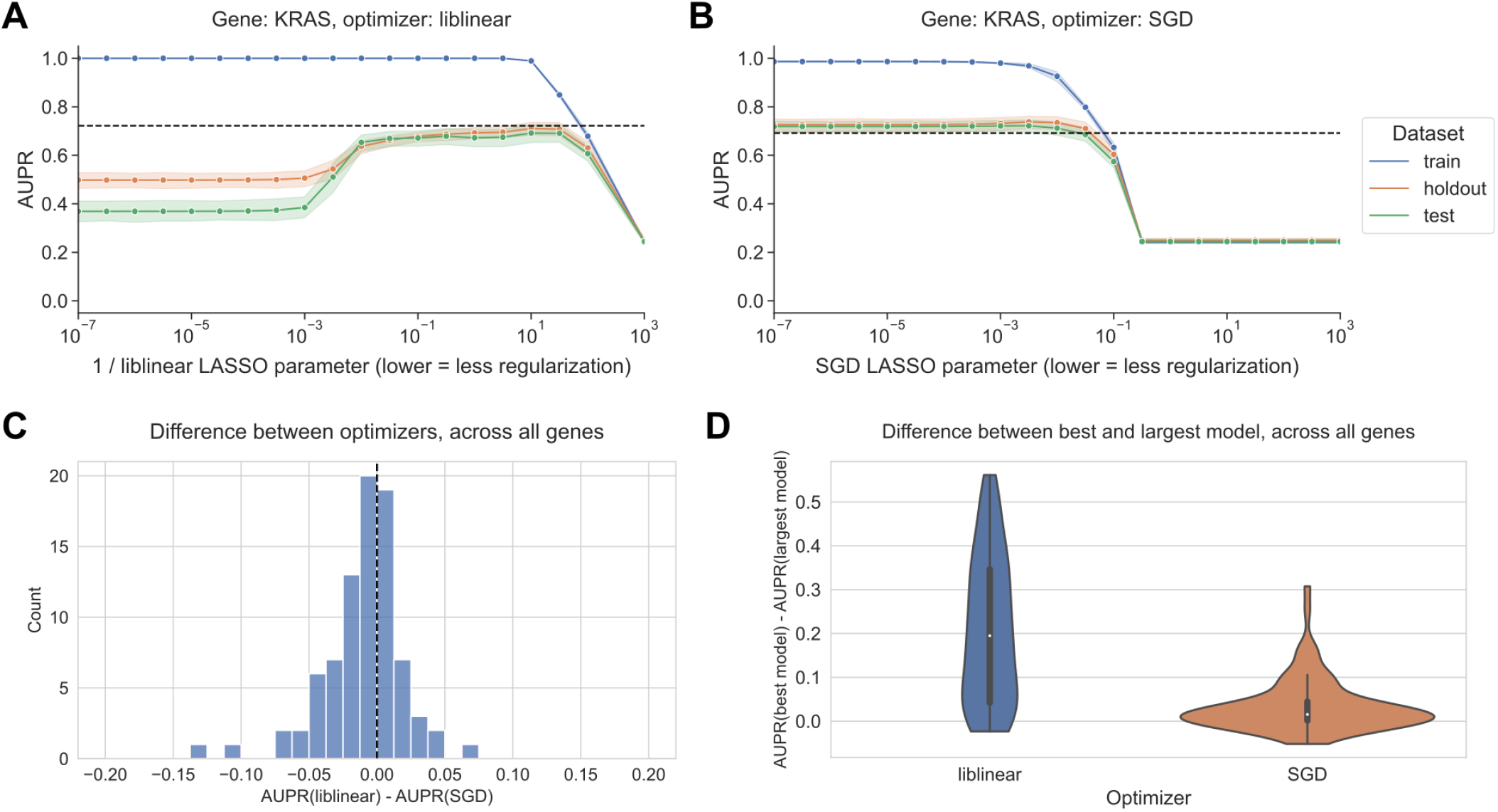
**A.** Performance vs. inverse regularization parameter for KRAS mutation status prediction, using the `liblinear` coordinate descent optimizer. Dotted lines indicate top performance value of the opposite optimizer. **B.** Performance vs. regularization parameter for KRAS mutation status prediction, using the SGD optimizer. “Holdout” dataset is used for SGD learning rate selection, “test” data is completely held out from model selection and used for evaluation. **C.** Distribution of performance difference between best-performing model for `liblinear` and SGD optimizers, across all 84 genes in Vogelstein driver gene set. Positive numbers on the x-axis indicate better performance using `liblinear`, and negative numbers indicate better performance using SGD. **D.** Distribution of performance difference between best-performing model and largest (least regularized) model, for `liblinear` and SGD, across all 84 genes. Smaller numbers on the y-axis indicate less overfitting, and larger numbers indicate more overfitting.

To determine how relative performance trends with `liblinear` tend to compare across the genes in the Vogelstein dataset at large, we looked at the difference in performance between optimizers for the best-performing models for each gene (Figure 1C). The distribution is centered around 0 and more or less symmetrical, suggesting that across the gene set, `liblinear` and SGD tend to perform comparably to one another. We saw that for 52/84 genes, performance for the best-performing model was better using SGD than `liblinear`, and for the other 32 genes performance was better using `liblinear`. In order to quantify whether the overfitting tendencies (or lack thereof) also hold across the gene set, we plotted the difference in performance between the best-performing model and the largest (least regularized) model; classifiers with a large difference in performance exhibit strong overfitting, and classifiers with a small difference in performance do not overfit (Figure 1D). For SGD, the least regularized models tend to perform comparably to the best-performing models, whereas for `liblinear` the distribution is wider suggesting that overfitting is more common.

### SGD is sensitive to learning rate selection

The SGD results shown in Figure 1 select the best-performing learning rate using a grid search on the holdout dataset, independently for each regularization parameter. We also compared against other learning rate scheduling approaches implemented in scikit-learn (see Methods for implementation details and grid search specifications). For KRAS mutation prediction, we observed that the choice of initial learning rate and scheduling approach affects performance significantly, and other approaches to selecting the learning rate performed poorly relative to `liblinear` (black dotted lines in Figure 2) and to the grid search. We did not observe an improvement in performance over `liblinear` or the grid search for learning rate schedulers that decrease across epochs (Figure 2A, C, and D), nor did we see comparable performance when we selected a single constant learning rate for all levels of regularization without the preceding grid search (Figure 2B). Notably, scikit-learn’s default “optimal” learning rate schedule performed relatively poorly for this problem, suggesting that tuning the learning rate and selecting a well-performing scheduler is a critical component of applying SGD successfully for this problem (Figure 2D). We observed similar trends across all genes in the Vogelstein gene set, with other learning rate scheduling approaches performing poorly in aggregate relative to both `liblinear` and SGD with the learning rate grid search (Supplementary Figure S2).

**Figure 2:**
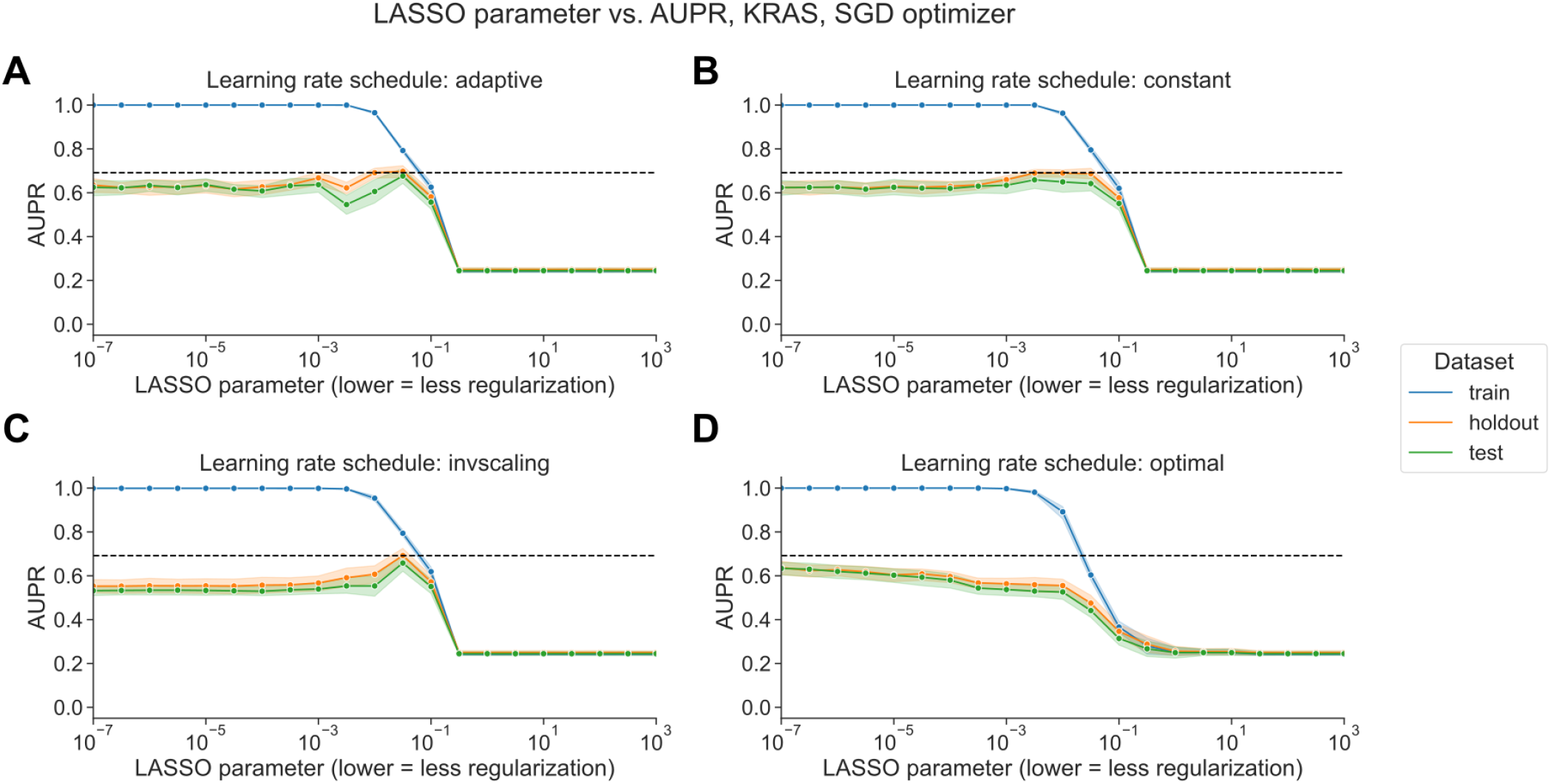
**A.** Performance vs. regularization parameter for KRAS mutation prediction, using SGD optimizer with adaptive learning rate scheduler. Dotted line indicates top performance value using `liblinear`, from Figure 1A. **B.** Performance vs. regularization parameter, using SGD optimizer with constant learning rate scheduler and a learning rate of 0.0005. **C.** Performance vs. regularization parameter, using SGD optimizer with inverse scaling learning rate scheduler. **D.** Performance vs. regularization parameter, using SGD optimizer with “optimal” learning rate scheduler.

### liblinear and SGD result in different models, with varying loss dynamics

We sought to determine whether there was a difference in the sparsity of the models resulting from the different optimization schemes. In general across all genes, the best-performing SGD models mostly tend to have many nonzero coefficients, but with a distinct positive tail, sometimes having few nonzero coefficients. By contrast, the `liblinear` models are generally sparser with fewer than 2500 nonzero coefficients, out of ∼16100 total input features, and a much narrower tail (Figure 3A). The sum of the coefficient magnitudes, however, tends to be smaller on average across all levels of regularization for SGD than for `liblinear` (Figure 3B). This effect is less pronounced for the other learning rate schedules shown in Figure 2, with the other options resulting in larger coefficient magnitudes (Supplementary Figure S3). These results suggest that the models fit by `liblinear` and SGD navigate the tradeoff between bias and variance in slightly different ways: `liblinear` tends to produce sparser models (more zero coefficients) as regularization increases, but if the learning rate is properly tuned, SGD coefficients tend to have smaller overall magnitudes as regularization increases.

**Figure 3:**
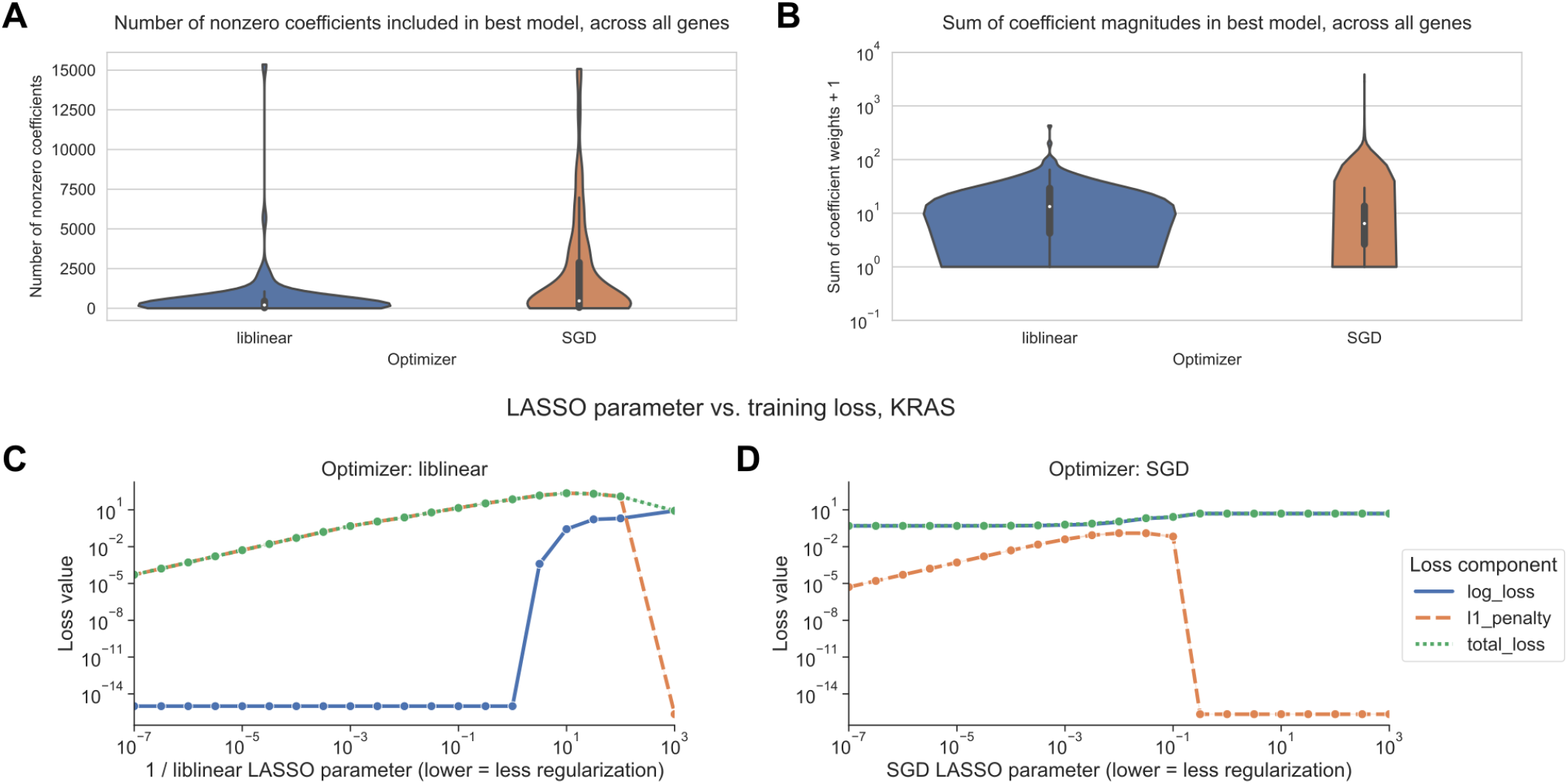
**A.** Distribution across genes of the number of nonzero coefficients included in best-performing LASSO logistic regression models. Violin plot density estimations are clipped at the ends of the observed data range, and boxes show the median/IQR. **B.** Distribution across genes of the sum of model coefficient weights for best-performing LASSO logistic regression models. **C.** Decomposition of loss function for models fit using `liblinear` across regularization levels. 0 values on the y-axis are rounded up to machine epsilon; i.e. 2.22 × 10^−16^. **D.** Decomposition of loss function for models fit using SGD across regularization levels. 0 values on the y-axis are rounded up to machine epsilon; i.e. 2.22 × 10^−16^.

We also compared the components of the loss function across different levels of regularization between optimizers. The LASSO logistic regression loss function can be broken down into a data-dependent component (the log-loss) and a parameter magnitude dependent component (the L1 penalty), which are added to get the total loss that is minimized by each optimizer; see Methods for additional details. As regularization strength decreases for `liblinear`, the data loss collapses to near 0, and the L1 penalty dominates the overall loss (Figure 3C). For SGD, on the other hand, the data loss decreases slightly as regularization strength decreases but remains relatively high (Figure 3D).

Other SGD learning rate schedules have similar loss curves to the `liblinear` results, although this does not result in improved classification performance (Supplementary Figure S4).

## Discussion

Our work shows that optimizer choice presents tradeoffs in model selection for cancer transcriptomics. We observed that LASSO logistic regression models for mutation status prediction fit using stochastic gradient descent were highly sensitive to learning rate tuning, but they tended to perform robustly across diverse levels of regularization and sparsity. Coordinate descent implemented in `liblinear` did not require learning rate tuning, but generally performed best for a narrow range of fairly sparse models, overfitting as regularization strength decreased. Tuning of regularization strength for `liblinear`, and learning rate (and regularization strength to a lesser degree) for SGD, are critical steps which must be considered as part of analysis pipelines. The sensitivity we observed to these details highlights the importance of reporting exactly what optimizer was used, and how the relevant hyperparameters were selected, in studies that use machine learning models for transcriptomic data.

To our knowledge, the phenomenon we observed with SGD has not been documented in other applications of machine learning to genomic or transcriptomic data. In recent years, however, the broader machine learning research community has identified and characterized implicit regularization for SGD in many settings, including overparametrized or feature-rich problems as is often the case in transcriptomics [14,15,16]. The resistance we observed of SGD-optimized models to decreased performance on held-out data as model complexity increases is often termed “benign overfitting”: overfit models, in the sense that they fit the training data perfectly and perform worse on the test data, can still outperform models that do not fit the training data as well or that have stronger explicit regularization. Benign overfitting has been attributed to optimization using SGD [16,17], and similar patterns have been observed for both linear models and deep neural networks [18,19].

Existing gene expression prediction benchmarks and pipelines typically use a single model implementation, and thus a single optimizer. We recommend thinking critically about optimizer choice, but this can be challenging for researchers that are inexperienced with machine learning or unfamiliar with how certain models are fit under the hood. For example, R’s `glmnet` package uses a cyclical coordinate descent algorithm to fit logistic regression models [20], which would presumably behave similarly to `liblinear`, but this is somewhat opaque in the `glmnet` documentation itself. Increased transparency and documentation in popular machine learning packages with respect to optimization, especially for models that are diffcult to fit or sensitive to hyperparameter settings, would benefit new and unfamiliar users.

Related to what we see in our SGD-optimized models, there exist other problems in gene expression analysis where using all available features is comparable to, or better than, using a subset. For example, using the full gene set improves correlations between preclinical cancer models and their tissue of origin, as compared to selecting genes based on variability or tissue-specificity [21]. On the other hand, when predicting cell line viability from gene expression profiles, selecting features by Pearson correlation improves performance over using all features, similar to our `liblinear` classifiers [22]. In future work, it could be useful to explore if the coefficients found by `liblinear` and SGD emphasize the same pathways or functional gene sets, or if there are patterns to which mutation status classifiers (or other cancer transcriptomics classifiers) perform better with more/fewer nonzero coefficients.

## Data and code availability

The data analyzed during this study were previously published as part of the TCGA Pan-Cancer Atlas project [23], and are available from the NIH NCI Genomic Data Commons (GDC). The scripts used to download and preprocess the datasets for this study are available at https://github.com/greenelab/pancancer-evaluation/tree/master/00_process_data, and the code used to carry out the analyses in this study is available at https://github.com/greenelab/pancancer-evaluation/tree/master/01_stratified_classification, both under the open-source BSD 3-clause license. Equivalent versions of Figure 1A and 1B for all 84 genes in the Vogelstein et al. 2013 gene set are available on Figshare at https://doi.org/10.6084/m9.figshare.22728644, under a CC0 license. This manuscript was written using Manubot [24] and is available on GitHub at https://github.com/greenelab/optimizer-manuscript under the CC0-1.0 license. This research was supported in part by the University of Pittsburgh Center for Research Computing through the resources provided. Specifically, this work used the HTC cluster, which is supported by NIH award number S10OD028483.

## Supplementary Material

**Figure S1:**
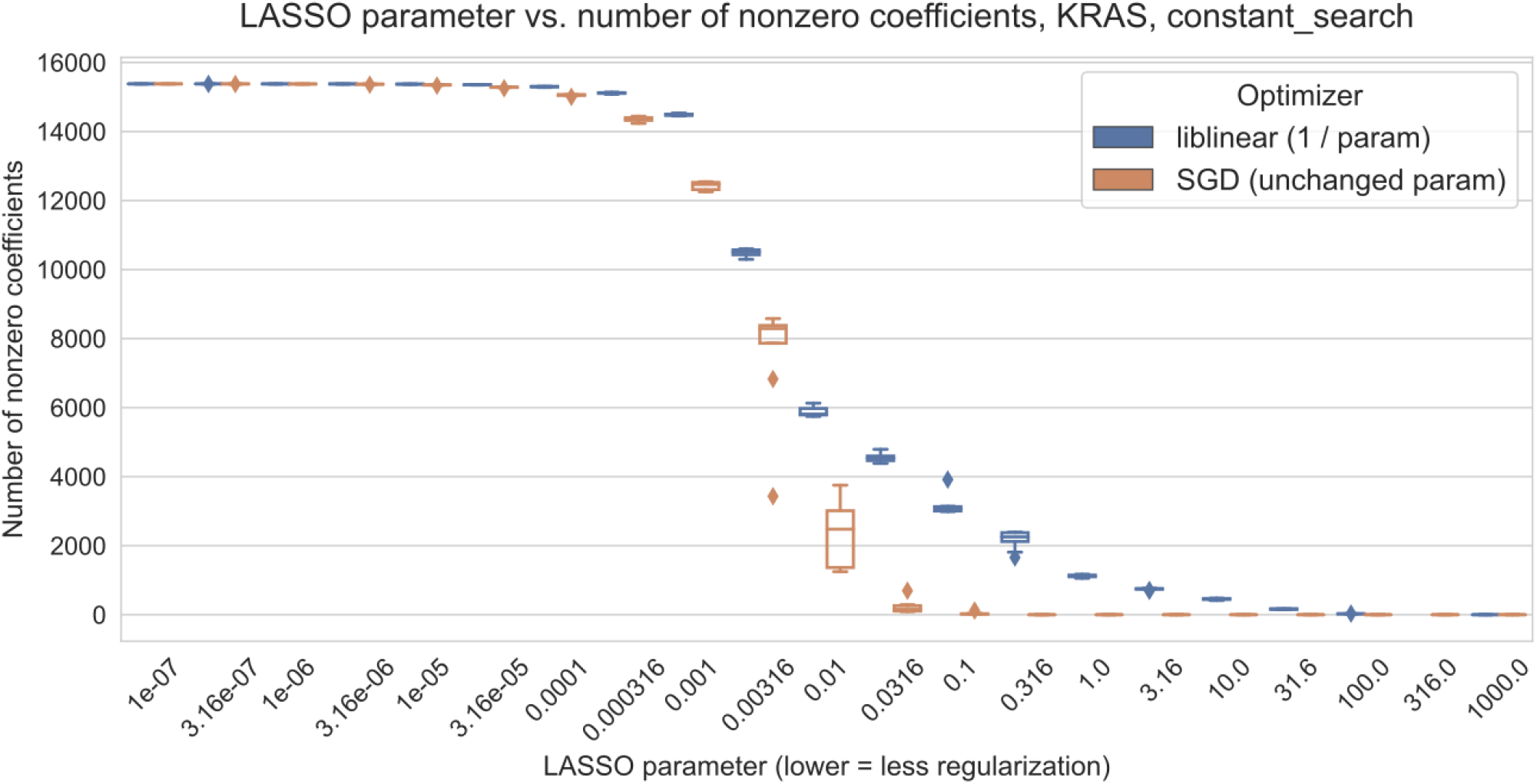
Number of nonzero coefficients (model sparsity) across varying regularization parameter settings for KRAS mutation prediction using SGD and `liblinear` optimizers.

**Figure S2:**
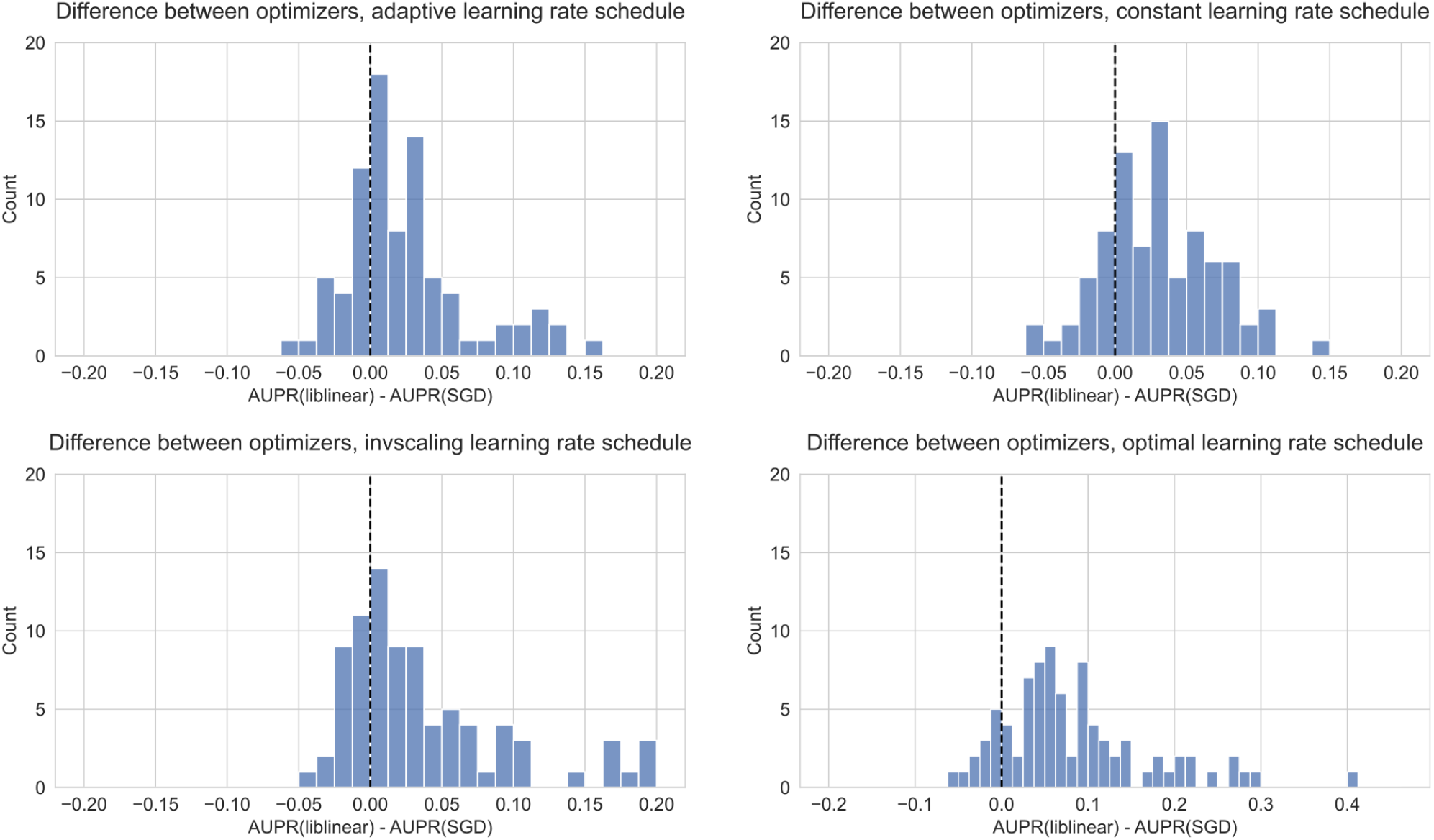
Distribution of performance difference between best-performing model for `liblinear` and SGD optimizers, across all 84 genes in Vogelstein driver gene set, for varying SGD learning rate schedulers. Positive numbers on the x-axis indicate better performance using `liblinear`, and negative numbers indicate better performance using SGD.

**Figure S3:**
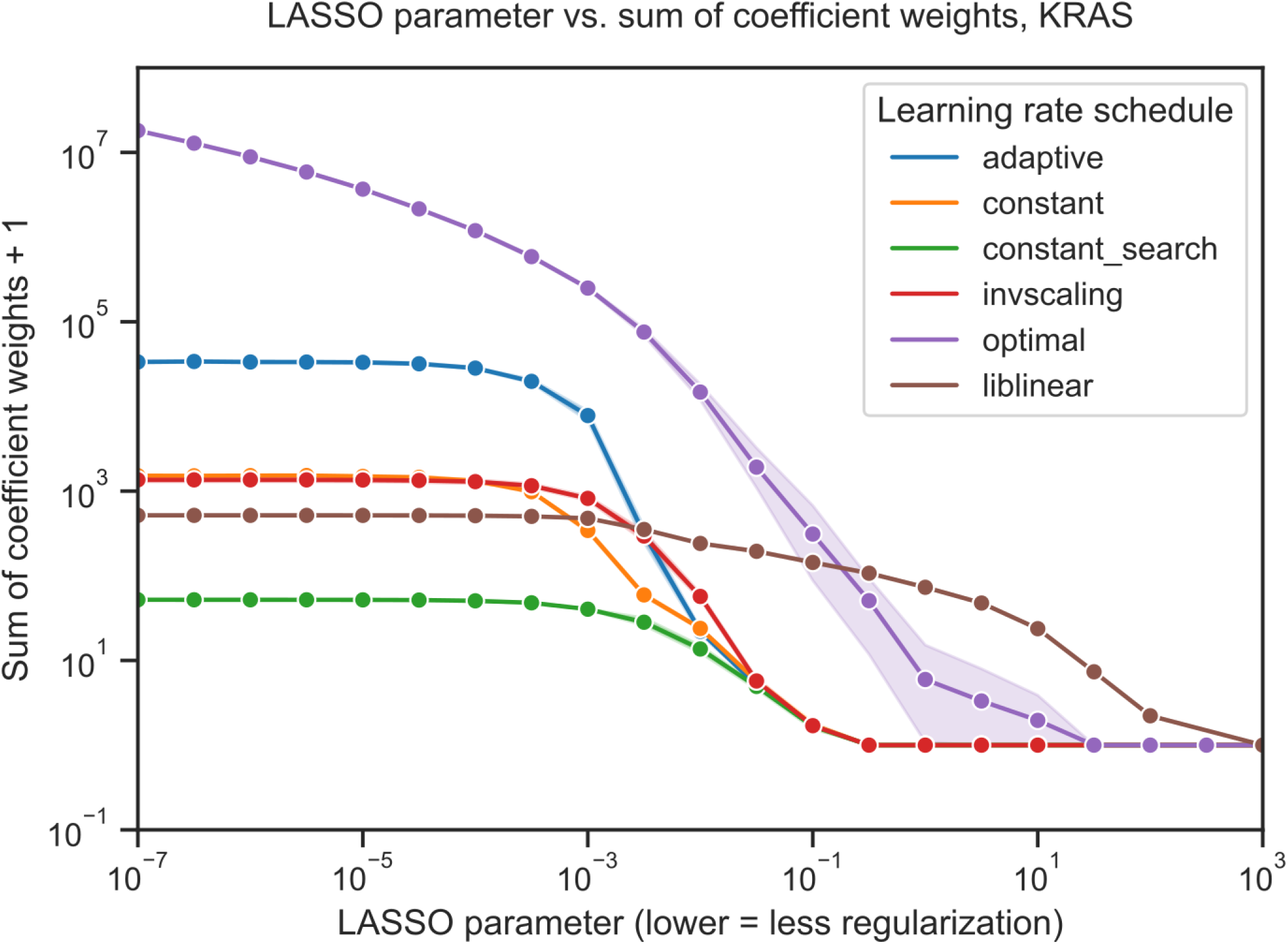
Sum of absolute value of coefficients + 1 for KRAS mutation prediction using SGD and `liblinear` optimizers, with varying learning rate schedules for SGD. Similar to the figures in the main paper, the `liblinear` x-axis represents the inverse of the *C* regularization parameter; SGD x-axes represent the untransformed *α* parameter.

**Figure S4:**
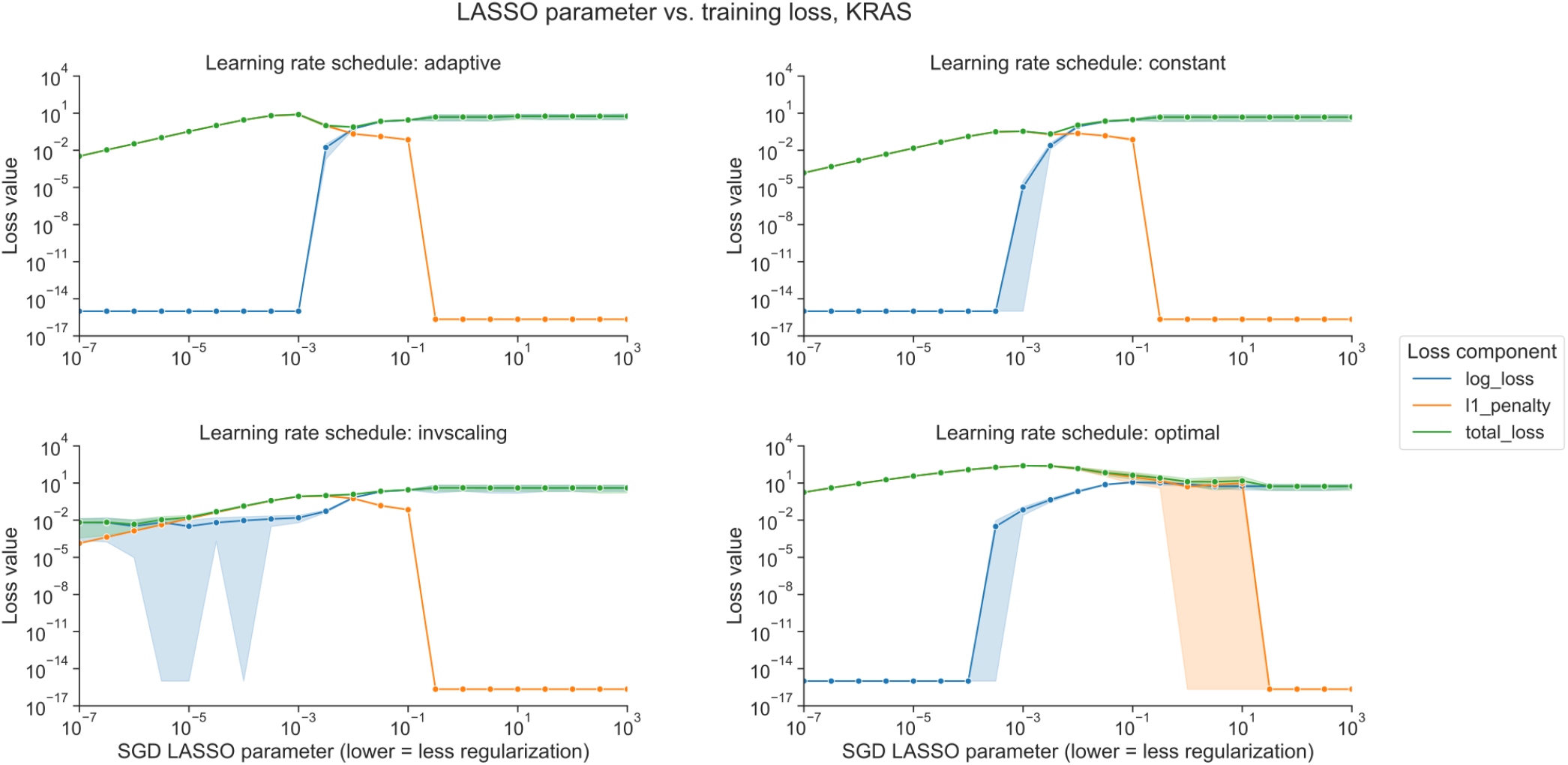
Decomposition of loss function into data loss and L1 penalty components for KRAS mutation prediction using SGD optimizer, across regularization levels, using varying learning rate schedulers. 0 values on the y-axis are rounded up to machine epsilon, i.e. 2.22 × 10^−16^.

## References

1. The ability to classify patients based on gene-expression data varies by algorithm and performance metric Stephen R Piccolo, Avery Mecham, Nathan P Golightly, Jérémie L Johnson, Dustin B Miller PLOS Computational Biology (2022-03-11) https://doi.org/gr43qd DOI: 10.1371/journal.pcbi.1009926 · PMID: 35275931 · PMCID: PMC8942277

2. Supervised learning is an accurate method for network-based gene classification Renming Liu, Christopher A Mancuso, Anna Yannakopoulos, Kayla A Johnson, Arjun Krishnan Bioinformatics (2020-04-14) https://doi.org/gmvnfc DOI: 10.1093/bioinformatics/btaa150 · PMID: 32129827 · PMCID: PMC7267831

3. Supervised Risk Predictor of Breast Cancer Based on Intrinsic Subtypes Joel S Parker, Michael Mullins, Maggie CU Cheang, Samuel Leung, David Voduc, Tammi Vickery, Sherri Davies, Christiane Fauron, Xiaping He, Zhiyuan Hu, … Philip S Bernard Journal of Clinical Oncology (2009-03-10) https://doi.org/c2688w DOI: 10.1200/jco.2008.18.1370 · PMID: 19204204 · PMCID: PMC2667820

4. Prediction of adjuvant chemotherapy benefit in endocrine responsive, early breast cancer using multigene assays Kathy S Albain, Soonmyung Paik, Laura van’t Veer The Breast (2009-10) https://doi.org/bp4rtw DOI: 10.1016/s0960-9776(09)70290-5 · PMID: 19914534

5. Scikit-learn: Machine Learning in Python Fabian Pedregosa, Gaël Varoquaux, Alexandre Gramfort, Vincent Michel, Bertrand Thirion, Olivier Grisel, Mathieu Blondel, Peter Prettenhofer, Ron Weiss, Vincent Dubourg, … Édouard Duchesnay Journal of Machine Learning Research (2011) http://jmlr.org/papers/v12/pedregosa11a.html

6. LIBLINEAR: A Library for Large Linear Classification Rong-En Fan, Kai-Wei Chang, Cho-Jui Hsieh, Xiang-Rui Wang, Chih-Jen Lin Journal of Machine Learning Research (2008) http://jmlr.org/papers/v9/fan08a.html

7. Online Learning and Stochastic Approximations Leon Bottou (1998) https://wiki.eecs.yorku.ca/course_archive/2012-13/F/6328/_media/bottou-onlinelearning-98.pdf

8. Cancer Genome Landscapes B Vogelstein, N Papadopoulos, VE Velculescu, S Zhou, LA Diaz, KW Kinzler Science (2013-03-28) https://doi.org/6rg DOI: 10.1126/science.1235122 · PMID: 23539594 · PMCID: PMC3749880

9. Scalable Open Science Approach for Mutation Calling of Tumor Exomes Using Multiple Genomic Pipelines Kyle Ellrott, Matthew H Bailey, Gordon Saksena, Kyle R Covington, Cyriac Kandoth, Chip Stewart, Julian Hess, Singer Ma, Kami E Chiotti, Michael McLellan, … Armaz Mariamidze Cell Systems (2018-03) https://doi.org/gf9twn DOI: 10.1016/j.cels.2018.03.002 · PMID: 29596782 · PMCID: PMC6075717

10. GISTIC2.0 facilitates sensitive and confident localization of the targets of focal somatic copy-number alteration in human cancers Craig H Mermel, Steven E Schumacher, Barbara Hill, Matthew L Meyerson, Rameen Beroukhim, Gad Getz Genome Biology (2011-04) https://doi.org/dzhjqh DOI: 10.1186/gb-2011-12-4-r41 · PMID: 21527027 · PMCID: PMC3218867

11. Widespread redundancy in -omics profiles of cancer mutation states Jake Crawford, Brock C Christensen, Maria Chikina, Casey S Greene Genome Biology (2022-06-27) https://doi.org/gqfqnm DOI: 10.1186/s13059-022-02705-y · PMID: 35761387 · PMCID: PMC9238138

12. Regression Shrinkage and Selection Via the Lasso Robert Tibshirani Journal of the Royal Statistical Society: Series B (Methodological) (1996-01) https://doi.org/gfn45m DOI: 10.1111/j.2517-6161.1996.tb02080.x

13. Machine Learning Detects Pan-cancer Ras Pathway Activation in The Cancer Genome Atlas Gregory P Way, Francisco Sanchez-Vega, Konnor La, Joshua Armenia, Walid K Chatila, Augustin Luna, Chris Sander, Andrew D Cherniack, Marco Mina, Giovanni Ciriello, … Armaz Mariamidze Cell Reports (2018-04) https://doi.org/gfspsb DOI: 10.1016/j.celrep.2018.03.046 · PMID: 29617658 · PMCID: PMC5918694

14. The Benefits of Implicit Regularization from SGD in Least Squares Problems Difan Zou, Jingfeng Wu, Vladimir Braverman, Quanquan Gu, Dean P Foster, Sham M Kakade arXiv (2022-07-12) https://arxiv.org/abs/2108.04552

15. Can Implicit Bias Explain Generalization? Stochastic Convex Optimization as a Case Study Assaf Dauber, Meir Feder, Tomer Koren, Roi Livni arXiv (2020-12-23) https://arxiv.org/abs/2003.06152

16. Benign Overfitting of Constant-Stepsize SGD for Linear Regression Difan Zou, Jingfeng Wu, Vladimir Braverman, Quanquan Gu, Sham Kakade Proceedings of Thirty Fourth Conference on Learning Theory (2021-07-21) https://proceedings.mlr.press/v134/zou21a.html

17. Understanding deep learning (still) requires rethinking generalization Chiyuan Zhang, Samy Bengio, Moritz Hardt, Benjamin Recht, Oriol Vinyals Communications of the ACM (2021-02-22) https://doi.org/gh57fd DOI: 10.1145/3446776

18. Benign overfitting in linear regression Peter L Bartlett, Philip M Long, Gábor Lugosi, Alexander Tsigler Proceedings of the National Academy of Sciences (2020-04-24) https://doi.org/gjgsxq DOI: 10.1073/pnas.1907378117 · PMID: 32332161 · PMCID: PMC7720150

19. Understanding deep learning requires rethinking generalization Chiyuan Zhang, Samy Bengio, Moritz Hardt, Benjamin Recht, Oriol Vinyals arXiv (2017-02-28) https://arxiv.org/abs/1611.03530

20. Regularization Paths for Generalized Linear Models via Coordinate Descent Jerome Friedman, Trevor Hastie, Robert Tibshirani Journal of Statistical Software (2010) https://doi.org/bb3d DOI: 10.18637/jss.v033.i01

21. Evaluating cancer cell line and patient-derived xenograft recapitulation of tumor and non-diseased tissue gene expression profiles<i>in silico</i> Avery S Williams, Elizabeth J Wilk, Jennifer L Fisher, Brittany N Lasseigne Cold Spring Harbor Laboratory (2023-04-13) https://doi.org/gr6jr4 DOI: 10.1101/2023.04.11.536431 · PMID: 37090499 · PMCID: PMC10120639

22. Gene expression has more power for predicting <i>in vitro</i> cancer cell vulnerabilities than genomics Joshua M Dempster, John M Krill-Burger, James M McFarland, Allison Warren, Jesse S Boehm, Francisca Vazquez, William C Hahn, Todd R Golub, Aviad Tsherniak Cold Spring Harbor Laboratory (2020-02-24) https://doi.org/ghczbr DOI: 10.1101/2020.02.21.959627

23. The Cancer Genome Atlas Pan-Cancer analysis project John N Weinstein, Eric A Collisson, Gordon B Mills, Kenna RMills Shaw, Brad A Ozenberger, Kyle Ellrott, Ilya Shmulevich, Chris Sander, Joshua M Stuart Nature Genetics (2013-09-26) https://doi.org/f3nt5c DOI: 10.1038/ng.2764 · PMID: 24071849 · PMCID: PMC3919969

24. Open collaborative writing with Manubot Daniel S Himmelstein, Vincent Rubinetti, David R Slochower, Dongbo Hu, Venkat S Malladi, Casey S Greene, Anthony Gitter PLOS Computational Biology (2019-06-24) https://doi.org/c7np DOI: 10.1371/journal.pcbi.1007128 · PMID: 31233491 · PMCID: PMC6611653

